# Magnetic resonance imaging for characterisation of a chick embryo model of cancer cell metastases

**DOI:** 10.1101/223891

**Authors:** Anne Herrmann, Arthur Taylor, Patricia Murray, Harish Poptani, Violaine Sée

**Affiliations:** Department of Biochemistry, University of Liverpool, Liverpool, L69 7ZB, UK; Centre for Preclinical Imaging, Department of Cellular and Molecular Physiology, University of Liverpool, Liverpool, L69 7ZB, UK

**Keywords:** MRI, chick embryo, imaging metastasis, neuroblastoma, *in ovo* imaging

## Abstract

**Background:** Metastasis is the most common cause of death for cancer patients, hence its study has rapidly expanded over the past few years. To fully understand all the steps involved in metastatic dissemination, *in vivo* models are required, of which murine ones are the most common. Therefore pre-clinical imaging methods have mainly been developed for small mammals. However, the potential of preclinical imaging techniques such as magnetic resonance imaging (MRI) to monitor cancer growth and metastasis in non-mammalian *in vivo* models is not commonly used. We have here used MRI to measure primary neuroblastoma tumour size and presence of metastatic dissemination in a chick embryo model. We compared its sensitivity and accuracy to end-point fluorescence detection.

**Methods:** Human neuroblastoma cells were labelled with GFP and micron-sized iron particles (MPIOs) and implanted on the extraembryonic chorioallantoic membrane of the chick embryo at E7. T2 RARE, T2 weighted FLASH as well as time-of-flight MR angiography imaging was applied at E14. Primary tumours as well as metastatic deposits in the chick embryo were dissected post imaging to compare with MRI results.

**Results:** MPIO labelling of neuroblastoma cells allowed *in ovo* observation of the primary tumour and tumour volume measurement non-invasively over time. Moreover, T_2_ weighted and FLASH imaging permitted the detection of very small metastatic deposits in the chick embryo.

**Conclusions:** The use of contrast agents enabled the detection of metastatic deposits of neuroblastoma cells in a chick embryo model, thereby reinforcing the potential of this cost efficient and convenient, 3R compliant, *in vivo* model for cancer research.

## BACKGROUND

Metastasis accounts for 90% of cancer deaths [1], yet it is one of the most poorly understood aspects of tumour progression. In order to reduce metastasis associated mortality, it is crucial to understand how, when and where metastasis occurs. However, small size, heterogeneity and large dispersal of disseminated cancer cells, combined with the limited sensitivity and spatial resolution of current clinical imaging methods, make their early and reliable detection challenging. Metastatic dissemination is a complex process involving several steps from the initial detachment of cells from the primary tumour, diffusion within the surrounding stromal tissue, degradation of the extracellular matrix and intravasation into the blood stream. Once in the circulatory system, tumour cells not only have to survive the hostile environment, but also attach to the endothelial cells of the vessel wall, extravasate in the extravascular tissue and proliferate in the metastatic site to form secondary tumours [2]. Although many of these steps have been studied at a molecular level *in vitro*, visualisation of the dynamic events *in vivo* remain elusive.

Currently used methods to detect the presence of metastasis *in vivo* in experimental studies rely mostly on endpoint measurements and require the termination of the experiment and organ dissection. Modern imaging modalities such as magnetic resonance imaging (MRI), positron emission tomography (PET) or bioluminescence imaging (BLI) allow non-invasive and longitudinal imaging of metastatic dissemination in whole organisms. In addition, MRI provides enhanced soft tissue contrast, three dimensional anatomical information and high spatial resolution. While the detection of primary tumours with MRI is already a routine practise, finding metastasis is more challenging as the metastatic cell population is heterogeneous and usually consists of single cells or a small group of malignant cells present in various tissue types, which makes their detection difficult. The use of contrast agents like iron oxide nanoparticles or Gadolinum-based agents for cell labelling can enhance contrast and thus detection limit. Iron oxide particles cause a distortion in the magnetic field leading to a change in T_2_/T_2_* relaxation and are mainly used to generate hypointense contrast on MRI [3, 4]. While a broad range of iron oxide particles are available for cell tracking, micron-sized iron particles (MPIOs) are of special importance as they are not only taken up efficiently and rapidly by cancer cells but also enable prolonged imaging due to their ability to label cells with a single particle only [5, 6]. Using contrast agents, metastasising cells could be detected in the lymph nodes [3, 7-9], liver [10-12] and brain [13] of rodents. Foster *et al*. reported the detection of approximately 100 MPIO labelled cells after direct implantation of melanoma cells in the lymph node [3]. Even detection at single cell level was observed as small metastastic deposits could be found in livers post mortem [11] and in the brain after injection into the left ventricle of the heart [13].

Whilst rodents constitute the most widely used model system for studying tumour development and metastasis, the chick embryo is a versatile 3R compliant model that is readily accessible *in* or *ex ovo*, nutritionally self-sufficient, cost-efficient and phylogenetically more similar to mammals than several other models of replacement, such as the zebrafish or nematode worm. The main advantage, when models for tumour formation are considered, is the accessibility of its chorioallantoic membrane (CAM), a highly vascularised extraembryonic membrane that is located directly beneath the eggshell. Thus, tumour cells can be engrafted easily, non-invasively and in the absence of an ‘interfering’ immune system, since the chick embryo is immunodeficient at earlier stages of development, when cells are implanted. Within days, tumour formation occurs and, in the case of aggressive tumours, metastasising cells can colonise the host’s organs via haematogenous metastasis [14]. However, despite all the advantages of the chick embryo, its potential has not been fully exploited so far.

We evaluated the advantages and limitations of MRI to study metastatic dissemination of neuroblastoma in the chick embryo. We have shown previously that we can induce metastasis *in vivo* by preculturing neuroblastoma cells in hypoxia or by treating with the hypoxia mimetic drug dimethyloxalylglycine (DMOG), where cells metastasise in 52% and 75% of cases, respectively. To understand the complex steps of the metastatic cascade and detect its onset, we investigated the feasibility of MRI to detect metastatic deposits of MPIO labelled neuroblastoma cells in the chick embryo.

## METHODS

### Cell culture

The human NB line SK-N-AS (ECACC No. 94092302) was grown in Minimal Essential Medium supplemented with 10% (v/v) foetal calf serum and 1% (v/v) non-essential amino acids (both Life Technologies) and maintained in a humidified incubator at 37°C, 5% CO_2_. For hypoxic studies, cells were maintained at 37°C, 5% CO_2_ and 1% O_2_ (Don Whitley Scientific – Hypoxystation-H35).

### Stable cell line generation and cell labelling

Lentiviral particles were produced with the transfer vector pLNT-SFFV-EGFP [15] as described previously [6]. For cell labelling 2 × 10^6^ SK-N-AS cells were seeded in a T-75 flask and allowed to grow for 24 h. Then 20 μM of Suncoast Yellow Encapsulated Magnetic Polymers (Bangs Beads, Stratech Scientific, Suffolk, England) were added directly to the complete culture medium and cells were allowed to grow for further 48 h. After the labelling period, the cells were carefully washed with phosphate buffered saline (PBS) to remove excess contrast agent, harvested and used for *in vivo* studies.

### Primary tumour, experimental and spontaneous metastasis assay

For the observation of the primary tumour, CAM implantation at E7 was performed as described previously [16]. In brief, fluorescent (GFP) and MPIO labelled SK-N-AS cells were harvested and 1 × 10^6^ cells/μl were resuspended in serum free media. CAM implantation was achieved by transferring 2 μl of the cell suspension into the CAM membrane fold created by careful laceration of White Leghorn chicken embryos.

For the observation of cells directly injected in the chick organs, fluorescent and MPIO labelled SK-N-AS cells were harvested and 1 × 10^5^ cells/μl resuspended in serum free media and 0.15% (v/v) Fast green (Sigma-Aldrich). Cell implantation was achieved by injecting 3 μl of the cell suspension into the brain of White Leghorn chicken embryos *in ovo* at E7 using a micro-capillary pipette.

For the observation of spontaneous metastasis CAM implantation at E7 was performed as described above for primary tumour formation using hypoxic preconditioned neuroblastoma cells. In brief, fluorescent (GFP) labelled SK-N-AS cells were preconditioned in 1% O_2_ for 3 days. MPIO labelling took place 48h prior harvesting. Cells were harvested and 1 × 10^6^ cells/μl were resuspended in serum free media. CAM implantation was achieved by transferring 2 μl of the cell suspension into the CAM membrane as explained above.

After cell implantation, eggs were maintained at 37 °C and 40% humidity until E11 or E14 and all animal work followed UK regulations (Consolidated version of ASPA 1986). For MR scanning, embryos were removed from the incubator at E11 or E14, cooled at 4 °C for 60 min and then imaged. In the case of time-of-flight (ToF) angiographic MRI, embryos were not cooled but anaesthetised with 3.6 mM Ketamine in 500 μl of PBS (Sigma-Aldrich) dropped directly onto the CAM prior to MR imaging.

### Fluorescent detection of tumour and metastatic deposits

Following MRI, a standard fluorescent stereo microscope (Leica M165-FC) was used to image primary tumours and metastatic deposits. Tumours were removed from the CAM and were imaged from three different perspectives (dorsal, ventral and lateral). Following removal of primary tumours from the CAM, embryos were dissected. Organs were removed and tumour cells and / or metastatic deposits identified by fluorescence.

Subsequently, tumour and organ samples were fixed for up to 12 h in 4% formaldehyde for the preparation of 10 μm thick frozen sections. Frozen tissue slices were stained with Hoechst and analysed with an epi-fluorescent microscope (Axio ObserverZ1, Zeiss). A representative sagittal MRI slice was correlated with the section of the region of tumour or metastatic deposit.

### Tumour volume calculation

#### By microscopy

Excised tumours were imaged from three different perspectives (dorsal, ventral and lateral). Average tumour volume was calculated as previously described [14] using 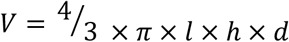, where l is length, h is height and d is depth.

#### By MRI

T_2_ weighted images were used for tumour volume calculation. The tumour area was measured with ImageJ (Wayne Rasband) in each slice and tumour volume was calculated using 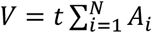, where N is the number of slices, Ai the area of the ROI encompassing the tumour and t the slice thickness [17].

### *In ovo* MRI

MR data were acquired with a Bruker Avance III spectrometer interfaced to a 9.4T magnet (Bruker Biospec 90/20 USR) using a 74 cm transmit-receive resonator coil. Sagittal images of the chick embryos were acquired using following sequences: **(1)** high resolution **TurboRARE** T2 weighted images with the following parameters: field of view 45 × 35 mm, matrix size 512 × 398 (256 × 198 for Figure 1A-C), slice thickness 0.4 mm (0.5 mm for Figure 1A-C), slice gap 0.3 mm, effective TE 35 ms, TR 7822 ms (6703 ms and 7262 ms for Figure 1A and Figure 1B/C, respectively), averages 5, slices 70 (60 and 65 for Figure 1A and Figure 1B/C, respectively), scan time 31 min 56 s (13 min 24 s and 14 min 31 s for Figure 1A and Figure 1B/C, respectively); **(2)** T2* weighted images using a **FLASH** sequence with the following parameters: field of view 45 × 35 mm, matrix size 512 × 398, slice thickness 0.4 mm, slice gap 0.3 mm, effective TE 6.88 ms, TR 1135 ms, averages 3, flip angle 30°, slices 70, scan time 22 min 35 s; **(3)** angiography using a **ToF** sequence with following parameters: field of view 45 × 35 mm, matrix size 512 × 398, slice thickness 0.4 mm, slice gap 0.3 mm, effective TE 3.1 ms, TR 13 ms, averages 2, flip angle 80°, slices 70, scan time 12 min 4 s.

**Figure 1.**
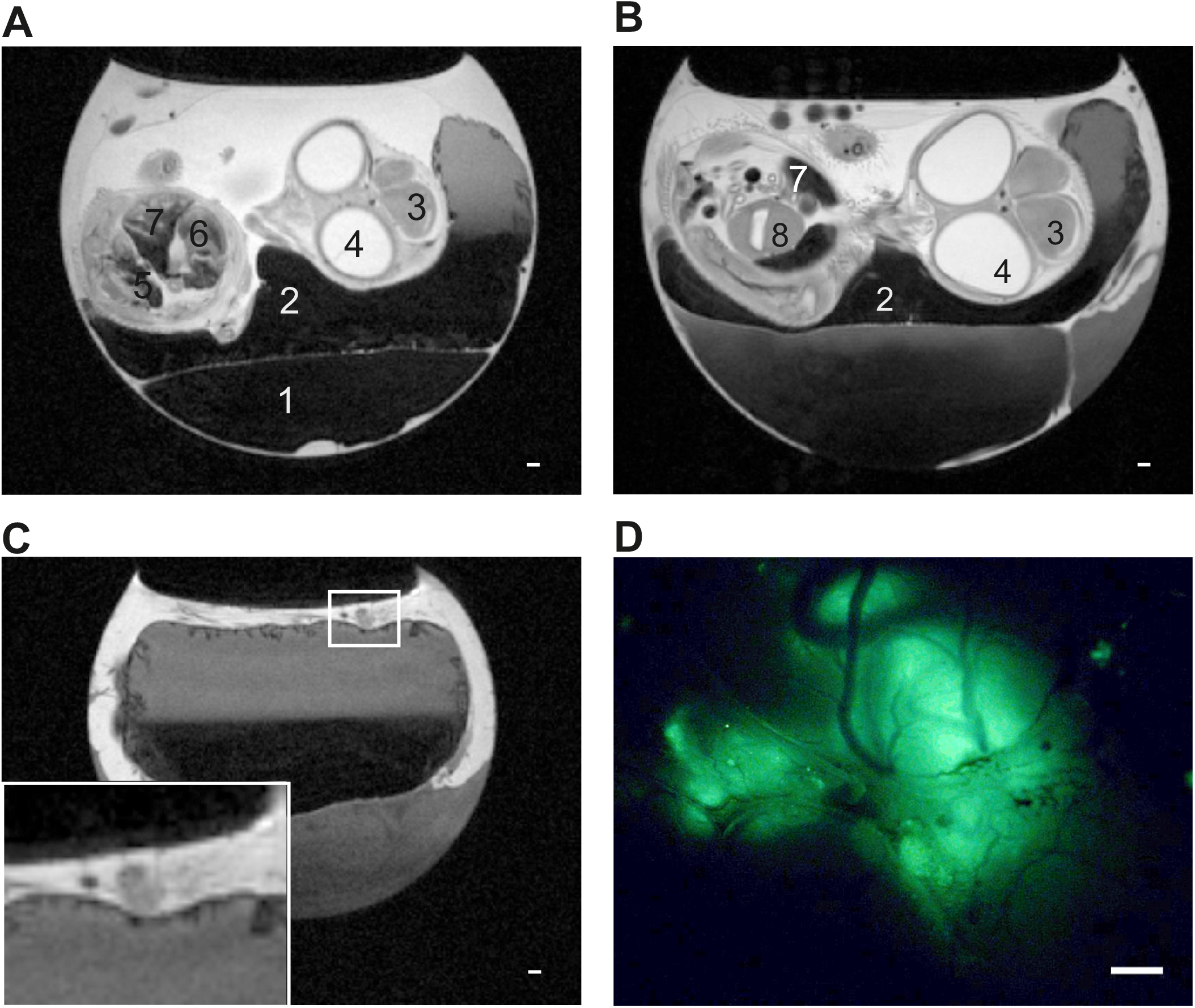
T2-weighted images of tumours growing on the CAM. **A and B)** Representative sagittal T_2_-weighted MR images of E11 (A) and E14 (B) chick embryo *in ovo*. Egg compartments like albumen (1), yolk (2) as well as chick embryo organs like brain (3),eyes (4), kidneys (5) heart (6) liver (7) and gizzard (8) can be identified. **C)** Representative sagittal T_2_-weighted MR images of embryonated chicken egg at E14 *in ovo*. Extraembryonic tumour can be identified on top of the CAM (zoom in inset) and correlates with fluorescent image (D). Due to the anatomy of the egg the primary tumour is not always located above the chick embryo and thus the chick embryo does not always appear in the same sagittal slice as the one showing the primary tumour. **D)** Representative fluorescence microscopy image of tumour formed by GFP-expressing neuroblastoma cells. Scale bars represents 1000 μm.

## RESULTS

### T_2_ weighted imaging of chick embryos allows observation of tumourigenesis and embryonic development

Fluorescently labelled (GFP) neuroblastoma cells were implanted on the CAM at E7 and tumour formation was assessed by MRI. Representative images from T_2_ weighted multi-slice MRI scans obtained at E11 and E14 are shown in Figure 1A and B, respectively. Allantois, yolk sack and chick embryo organs such as liver, kidneys and heart can be clearly identified and studied over time. The cooling of the embryos at 4 °C for 60 min prior to imaging reduced their movement for up to 60 min allowing motionless imaging. Tumours grown on the CAM can be easily identified by MRI (Figure 1C). Primary tumour dissection with a fluorescent microscope revealed that location and morphology of the tumour are in good correlation with the images acquired by MRI (Figure 1D).

### MPIO labelling facilitates tumorigenesis observation and allows tumour volume measurement

To investigate whether MPIO labelling enhances the detection of primary tumours, GFP-expressing neuroblastoma cells were labelled with red fluorescent MPIOs for 48 h prior to CAM implantation. MPIO uptake was efficient as all cells contained multiple MPIOs 24 h post labelling (Figure 2A). MPIO labelled cells successfully formed tumours on the CAM and signal from GFP as well as MPIOs could be detected by fluorescence (Figure 2B). Tumour formation was then assessed by T_2_ weighted and T_2_* fast low angle shot (FLASH) MRI scans. Using FLASH, areas containing cells labelled with MPIOs should experience an enhanced signal loss compared to other areas, such as blood vessels or tissue. Representative images from T_2_ weighted and T_2_* FLASH scans obtained at E14 show that, as with unlabelled cells, tumours could be readily identified in the MR scans (Figure 2C). Tumours formed from MPIO labelled cells, however, displayed a much stronger signal loss, which is expected given their iron oxide load. Primary tumour dissection revealed that location and morphology of the tumour were comparable to the images acquired by MRI (Figure 2B). Fluorescent images of frozen tumour sections revealed a homogenous distribution of MPIOs within the tumour (Figure 2D). Only a fraction of cells still contained MPIOs, which was expected due to extensive cell proliferation during tumour development *in vivo* and consequently, progressive dilution of the label between daughter cells. MPIOs were only observed in GFP-labelled tumour cells and not in the surrounding chick tissue.

**Figure 2.**
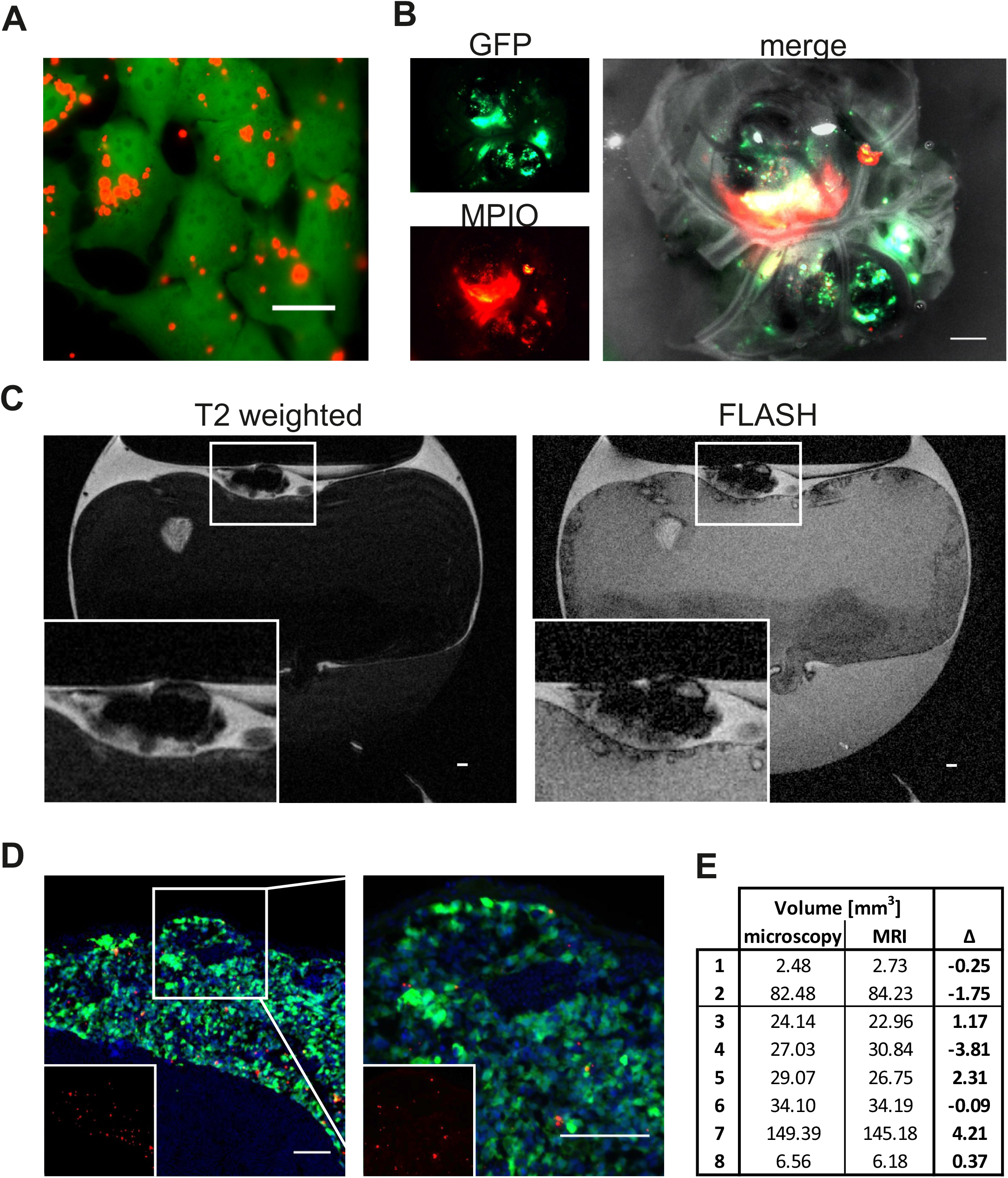
T2-weighted and T2*FLASH images of tumours labelled with MPIO. **A)** GFP-expressing SK-N-AS cells (green) 24 h post labelling with 20 μM MPIO (Suncoast Yellow Encapsulated Magnetic Polymers - Bangs Beads, red). Scale bar is 20 μm. **B)** Single channel and overlay image of neuroblastoma tumour post dissection formed by GFP-expressing SK-N-AS cells (green) which were labelled with MPIO (red) 48h prior CAM implantation. Scale bar is 1000 μm. **C)** Representative sagittal T_2_-weighted and T_2_* FLASH MR images of E14 (A) chick embryo *in ovo*. Tumour formed by cells labelled with MPIO can be identified on top of the CAM (zoom in inset). Scale bar is 1000 μm. **D)** Representative image of tumour formed on the CAM by GFP-expressing SK-N-AS cells (green) labelled with MPIO (red). Nuclei are stained with Hoechst (blue). Inset shows MPIO only (red). Right image is 2.5xzoom. Scale bar is 100 μm. **E)** Comparison of tumour volume [mm^3^] measured by microscopy or MRI. Tumours 1–2 were formed by cells without MPIO, tumours 3–8 were formed by cells with MPIO.

In order to determine whether MRI can also be used to determine tumour volume, tumour areas on sagittal T_2_ weighted MRI slices displaying the primary tumour were measured and calculated as described in the method section. Tumour volume estimates were then compared to those obtained from tumour excision and microscopy and were comparable (Figure 2E). The slight dissimilarity between the two methods can be explained by the difference in volume calculation. While the volume measured in images obtained by microscopy assumes that the tumour is a spherical object, MRI allows a more precise estimation as the area of each slice displaying a tumour is considered. Whilst it was easier to see the tumours when they were labelled with MPIO, it did not drastically change the ability to detect primary tumours and it had no impact on tumour volume measurements. Thus, MRI can easily be used to study the presence, progression and volume of tumours non-invasively over time, in contrast to fluorescence microscopy, which necessitate tumour excision from the CAM.

### MPIO labelling combined with T_2_ weighted and FLASH imaging allows detection of metastasis

To first investigate whether MPIO labelling enables the detection of cells within the chick embryo organs, 3 × 10^5^ GFP-expressing and MPIO labelled neuroblastoma cells were directly injected into the brain of the chick embryo at E7 and analysed at E14. Representative images from T_2_ weighted and T_2_* FLASH scans obtained at E14 are shown in Figure 3A. A small region (2 × 1 mm) of signal loss can be observed in the brain indicating the presence of MPIO labelled tumour cells. Size, shape and location of the cell cluster correlate well with the fluorescent signal obtained by subsequent fluorescence microscopy and tissue analysis (Figure 3B). Like in the primary tumour growing on the CAM, MPIOs were homogenously distributed among the cell population, with a great proportion of the cells not containing MPIOs anymore.

**Figure 3.**
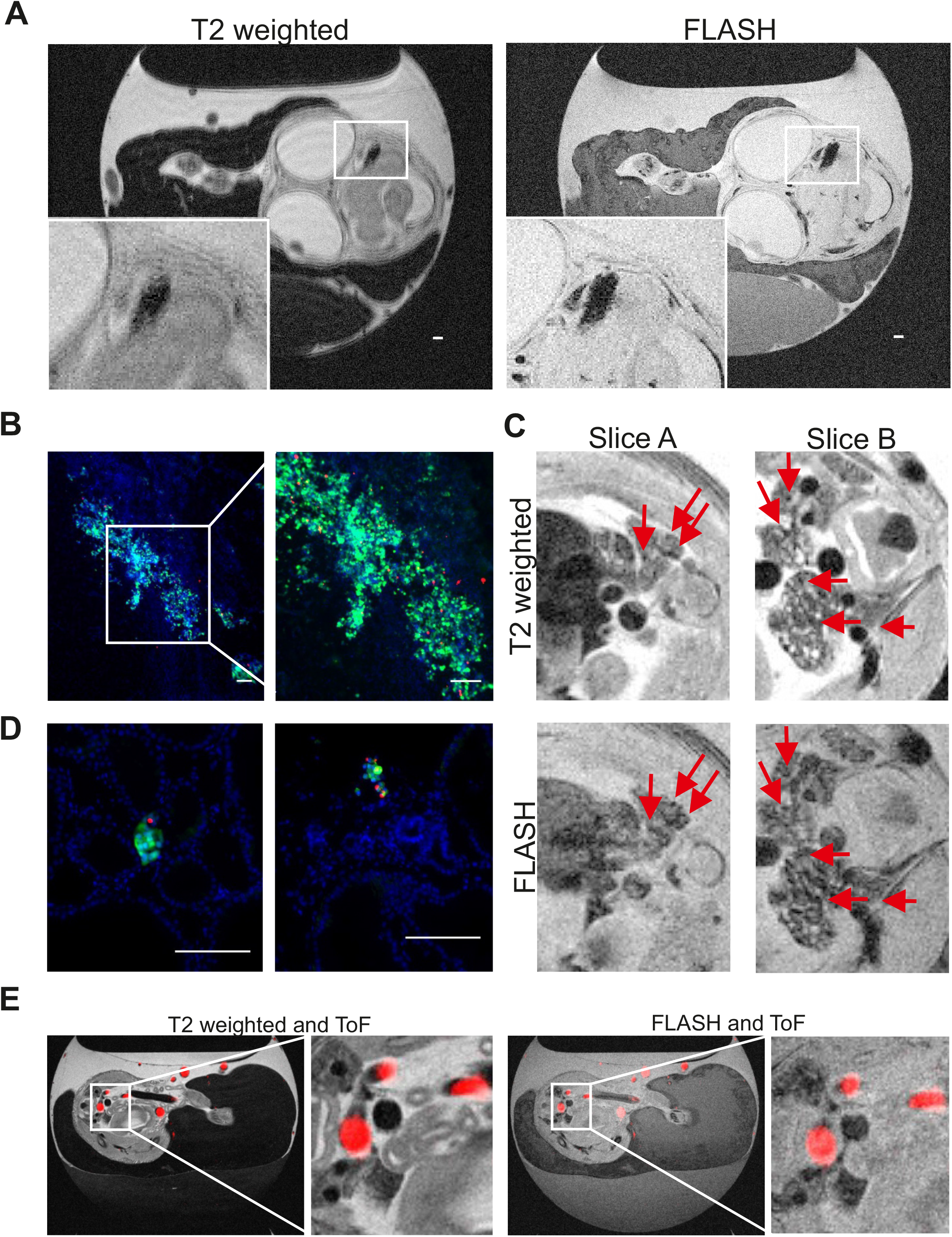
MR images of cell deposits and metastasis in the chick embryo organs. **A)** Representative sagittal T_2_-weighted and T_2_* FLASH MR images of E14 chick embryo *in ovo*. Metastatic deposit formed by cells labelled with MPIO can be identified in the brain (zoom in inset) Scale bar is 1000 μm. **B)** Representative fluorescence microscopy image of brain slice showing cluster of GFP and MPIO labelled neuroblastoma cells, and zoom. Scale bar is 100 μm. C) Representative sagittal T_2_-weighted and T_2_* FLASH MR images of E14 chick embryo *in ovo*. Shown are two slices of the abdominal region and kidneys. Arrows indicate signal loss that intensified in the T_2_* FLASH sequence and thus indicates the potential presence of metastasis. **D)** Representative fluorescence microscopy image of kidney slices showing metastatic deposit of GFP and MPIO labelled neuroblastoma cells. Scale bar is 100 μm. **E)** Representative sagittal T_2_-weighted and T_2_* FLASH MR images of E14 chick embryo *in ovo* combined with ToF MRA (red).

We further evaluated whether MPIO labelling enables the detection of spontaneous and smaller metastasis by using a spontaneous metastasis model in the chick embryo [14]. We have previously shown that we can control metastasis of neuroblastoma cells by hypoxic preconditioning [14]. While cells grown in normoxia are capable of tumourigenesis but not of metastatic invasion, cells grown in hypoxia (3 days in 1% O_2_) metastasise in 52% of the cases from the primary tumour into the chick embryo organs. Whilst such metastatic phenotype was observed as an end point measurement upon chick organs dissection, the detection of metastasis in the chick embryo using imaging modalities has not yet been reported. GFP-expressing and MPIO labelled neuroblastoma cells were cultured under hypoxia, implanted on the CAM at E7 and their metastasis into chick tissues was assessed at E14, using T_2_ weighted and T_2_* FLASH scans (Figure 3C). FLASH MRI was applied in order to distinguish the regions of signal loss caused by small blood vessels, haemorrhagic areas, air-tissue interfaces like the pancreas or areas devoid of proton signal such as the lungs from potential neuroblastoma metastasis. Several small areas of signal loss were observed in the kidneys of chick embryos. Arrows indicate the areas where signal loss with T_2_* FLASH was maintained or increased, indicating the presence of metastasising labelled cells. Organ dissection and analysis by fluorescence microscopy confirmed the presence of several metastatic deposits in the kidney as shown in Figure 3D. The metastatic deposits consisted of up to 12 cells and up to 4 MPIOs. Thus even very small metastasis could be detected by MRI. However, their identification was not trivial given their small size in the inherent low MR signal of the kidney.

To be able to differentiate the small metastatic deposits from the blood vessels, we applied time-of-flight MR angiography (ToF MRA). This allowed signal loss caused by small blood vessels to be distinguished from potential metastasis more effectively than using FLASH alone. As ToF is dependent on the influx of fresh unsaturated blood, chick movement reduction was necessary. We tested two methods for reducing embryo movement: ketamine anaesthesia and embryo cooling. Cooling the embryos resulted in a reduced blood flow making successful ToF acquisition unfeasible. Therefore, ketamine anaesthesia was used. Representative ToF images were overlaid with imaged from T_2_ weighted and T_2_* FLASH scans obtained at E14 are shown in Figure 3E. Although ToF MRA allows the detection of bigger blood vessels, the small and very fine vessels in the kidney for example as well as other hypointense areas such as the gastrointestinal tract could not be resolved with the current acquisition protocol that was optimized to keep the embryos viable limiting the ability to detect small metastasis in this model.

Taken together, we have demonstrated that primary tumour formation on the CAM can be easily detected in the chick embryo model. Tumour cells within the organs of the chick embryo can be also detected, however 12 cells (labelled with 1 remaining MPIO) seemed to constitute the lower limit for a reliable detection and we anticipate that larger metastases are required to provide a more robust signal.

## DISCUSSION

Much of our current understanding about the complex metastatic process comes from modern imaging techniques. While each imaging modality comes with advantages and limitations, MRI offers detailed three dimensional anatomical information and high resolution over time in a non-invasive manner. In agreement with others, we show here, that MRI is a powerful imaging modality for the study of tumour progression [18, 19] and embryonic development [20-24] in the chick embryo. Compared to optical imaging, it can be used to detect the presence of tumours even when they are hidden beneath the egg shell and allows the non-invasive study of tumour progression and volume over time.

The main aim of this study was to determine whether MRI can also be used to detect the presence of metastasis non-invasively in the chick embryo. In order to observe metastatic dissemination, neuroblastoma cells were labelled with MPIOs as contrast agents. The labelling of cancer cells with MPIOs did not alter tumour formation on the CAM. While it did not offer significant advantages for primary tumour detection compared to unlabelled cells, it was necessary for small metastasis detection in the chick embryo organs. We initially tried to detect large clusters of cells, administered directly to the brain of the chick embryo, which resulted in a substantial loss of signal and thus in a robust detection of cancer cells. This suggests that cancer lesions of about 2 mm are detectable in the chick embryo, a size that is smaller than the current MRI detection limit of metastasis of 10–20 mm in rodents [25]. This finding is in agreement with others that have used SPIONs to successfully detect micrometastases in lung, lymph node and brain in mice [3, 13, 26]. Using SPIONs and hyperpolarized ^3^He MRI, Branca *et al*. could detect micrometastasis of 0.3 mm in the lung of mice [26], while Foster *et al*. could detect 100 MPIO labelled cells by MRI after injecting them directly into the lymph node of mice [3] and Heyn *et al*. used SPION labelled breast cancer cells to detect a small number of cells in the brain of mice [13]. We have also shown previously that MRI can be used to reliably detect cell clusters of 5 × 10^4^ SPION labelled cells in the brain *ex vivo* [6]. Apart from SPIONs, other contrast agents have also been used for imaging metastasis. In mice, Zhou *et al*. could detect breast cancer metastases of less than 0.5 mm in different organs such as the lung, liver, lymph node, adrenal gland and bone using a gadolinium based contrast agent [27]. Xue *et al*. developed a protein-based contrast agent that enabled them to image early liver metastases as small as 0.24 mm in diameter after tail vein injection of uveal melanoma cells into mice [25].

Compared to mice, the chick embryo is a cost effective and convenient model, complying with the 3Rs by replacement of animal use. At E14, the metastatic deposits of neuroblastoma cells consist of only few cells, hence we wanted to determine if MRI could be used for their identification. We could observe signal reduction caused by a single MPIO particle and thus identify very small clusters of metastasised cells in the kidneys. However, the signal observed in the chick’s internal organs, including the kidney is inherently low, hence reliable detection of metastatic cells remains challenging with a potential of increased false positives. A confirmatory method, such as dissection, was needed to confirm the presence of such small metastatic deposits. These detection difficulties can be partially overcome by applying special techniques, such as ToF MRA, which we used here. Although ToF MRA enabled us to identify larger blood vessels, very small blood vessels couldn’t be resolved and consequently failed to facilitate the reliable detection of small metastasis in organs such as the kidney. Thus, the detection of metastasis of tumour types that disseminate either in organs of minimal signal loss, such as the brain or disseminate in bigger cell clusters, is more appropriate to this model.

In conclusion, we report that MRI is a suitable and highly sensitive imaging modality to image tumourigenesis *in vivo* using a chick embryo. We could, for the first time, identify metastatic deposits in the chick embryo by MRI. However, for reliable detection, we observed that 12 cells was the lower limit of detection. Whilst this means that this approach cannot be used to detect the onset of metastasis from a single cell, the small metastases observed was still remarkable, with the potential of providing longitudinal view of disease progression in the same animal non-invasively, particularly of primary tumours generated in areas such as the CAM or injected cells in the brain.

## LIST OF ABBREVIATIONS

BLI: bioluminescence imaging
CAM: chorioallantoic membrane
DMOG: Dimethyloxalylglycine
FLASH: fast low angle shot
H&E: haematoxylin and eosin
MPIOs: micron-sized iron particles
MRI: magnetic resonance imaging
PBS: phosphate buffered saline
PET: positron emission tomography
ToF MRA: time-of-flight MR angiography

## DECLARATIONS

### Ethics approval and consent to participate

Not applicable, as the experiments with chick embryos were terminated at E14 and hence the proposed model is classified as non-protected under the Animals Scientific Procedures Act 1986 (amended 2012).

### Consent for publication

Not applicable.

### Availability of data and materials

The full MRI scans and images are all available in figshare https://doi.org/10.6084/m9.figshare.5625271.v1

Figure 1A shows slice 45 from T2-weighted MR Scan ‘Fig1A’ in ovo at E11. Figure 1B shows slice 44 from T2-weighted MR Scan ‘Fig1B-C’ in ovo at E14. Figure 1C shows slice 12 from T2-weighted MR Scan ‘Fig1B-C’ in ovo at E14. Figure 2C shows slice 23 from T2-weighted and T2* FLASH MR Scan ‘Fig2C T2’ and ‘Fig2C FLASH’ in ovo at E14. Figure 3A shows slice 14 from T2-weighted and T2* FLASH MR Scan ‘Fig3A T2’ and ‘Fig3A FLASH’ in ovo at E14. Figure 3C shows slice 34 and 42 from T2-weighted and T2* FLASH MR Scan ‘Fig3C T2’ and ‘Fig3C FLASH’ in ovo at E14. Figure 3E shows slice 35 from T2-weighted and T2* FLASH MR Scan ‘Fig3E T2 ToF’ and ‘Fig3E FLASH_ToF’ in ovo at E14.

### Authors' contributions

A.H. and A.T. performed the experiments, analysed the data, prepared figures and drafted the manuscript. V.S. conceived and coordinated the study and wrote the manuscript. T.M, D.M. and H.P. contributed to the direction of the project and provided input in data analysis. All authors reviewed drafts of the manuscript and gave final approval for publication.

## Acknowledgements

Imaging data in this article were obtained in the Centre for Cell Imaging (CCI) and in the Centre for Preclinical Imaging (CPI) of the University of Liverpool. The equipment used in the CCI and the CPI has been funded by the Medical Research Council (MRC) (MR/K015931/1 and MR/L012707/1, respectively).

